# Mapping bacterial cutinase sequence space by high-throughput screening reveals that PET hydrolysis is a rare property

**DOI:** 10.64898/2026.08.20.745939

**Authors:** Robin Dorau, Malene Billeskov Keller, Emma Molter Thiesen, Johanna Katarina Sofie Tiemann, Morten Gjermansen, Pengfei Tian, Kim Borch, Kenneth Jensen, Peter Westh

## Abstract

Poly(ethylene terephthalate) (PET) is one of the most widely produced plastics, and enzymatic depolymerization offers a promising route to closed-loop recycling under mild conditions. However, most known bacterial PET hydrolases belong to a conserved canonical-fold cutinase family, leaving much of α/β-hydrolase diversity unexplored. Here, we mapped bacterial cutinase sequence space by combining bioinformatics-guided sequence selection with high-throughput secretion screening in *Bacillus subtilis*. A library of 1,120 genes encoding 954 unique bacterial cutinases, spanning canonical- and minimal-fold families, was screened for activity on Impranil DLN and semicrystalline PET. We identified 156 secreted cutinases with polyester activity, broadly distributed across sequence space, but only ten showed detectable PET hydrolysis, all from the canonical-fold family. These PET hydrolases were active at 40-50°C, preferred alkaline pH, and showed moderate thermostability. Our results demonstrate that PET activity is rare among bacterial cutinases and provide a scalable workflow for discovering diverse enzyme starting points.

**Significance Statement:** Poly(ethylene terephthalate) (PET) is a major plastic waste stream, and enzymatic depolymerization offers a route to closed-loop recycling under mild conditions. However, known PET-degrading enzymes occupy only a narrow region of bacterial cutinase sequence space, limiting discovery of new enzyme scaffolds. We combined bioinformatics-guided sequence selection with high-throughput screening in an industrial host to survey more than 1,100 bacterial cutinase genes. This approach identified 156 secreted polyester-active enzymes, including ten previously unreported PET hydrolases. This work provides an experimentally grounded map of polyester and PET hydrolysis across bacterial cutinase diversity, revealing that PET-degrading activity is relatively rare despite widespread polyester hydrolysis and expanding the foundation for future enzyme discovery and engineering.

## Introduction

In recent years, biocatalysis has emerged as a promising strategy for the recycling of ester-based plastics, particularly poly(ethylene terephthalate) (PET), one of the most abundantly produced synthetic polymers (1, 2) . PET hydrolases have attracted significant attention because they can selectively depolymerize PET into its constituent monomers under relatively mild reaction conditions (3, 4). This approach provides a more sustainable alternative to classical mechanical recycling, facilitating efficient monomer recovery and the re-synthesis of virgin PET. Such closed-loop recycling is central to advancing a circular economy for plastics (4–7).

Several naturally occurring hydrolase enzymes with PET-degrading activity have been identified, most of which exhibit promiscuous activity related to their native function as cutinases, lipases, or carboxylesterases (8, 9). As of this writing, the Plastics-Active Enzymes (PAZy) Database lists 311 PET hydrolases (10). The most efficient and extensively studied enzymes are derived from bacteria or metagenomes and all belong to the α/β-hydrolase family PF12695 (InterPro: IPR029059), which are built from eight β-sheets flanked by α-helices representing the canonical α/β-hydrolase-fold described by Ollis et al. (11). Enzymes from this family are referred to here as canonical-fold cutinases. Despite the growing number of PET-active enzymes in this family, their structural and sequence diversity remains highly conserved, reflecting their close phylogenetic relationships (12).

In contrast, a smaller number of fungal PET hydrolases has been reported, with the most efficient examples belonging to the cutinase/acetylxylan esterase family PF01083 (InterPro: IPR000675), referred to here as minimal-fold cutinases. These enzymes form a distinct phylogenetic branch, and while they also belong to the α/β-hydrolase superfamily, they are typically smaller than canonical-fold cutinases with only five β-sheets flanked by α-helices. Bacterial enzymes belonging to the minimal-fold cutinases are also known, but so far only two have been demonstrated to act on plastics (13, 14).

The discovery of new PET-active enzymes is often limited to computational mining efforts, as experimental exploration of large sequence spaces remains constrained by limited experimental throughput allowing for application-relevant conclusions and limitations of traditional gene synthesis techniques. Conventional phosphoramidite chemistry restricts oligonucleotide synthesis to fewer than three hundred bases, which necessitates additional assembly steps to obtain full-length genes. Although synthetic DNA is now routinely used for gene cloning, the excessive cost of the fragments remains a major limitation. Therefore, experimental efforts regarding PET hydrolases have typically been limited in scale, including analyses with up to 475 genes in the study by Norton-Baker et al. and 232 genes by Seo et al. (15, 16). Synthesis approaches to overcome those limitations have been developed, such as the Dropsynth approach (17–19), but have not yet been applied for exploring the sequence space of α/β-hydrolases to discover phylogenetically distant enzymes with activity against plastics.

In this study, we experimentally explored the bacterial cutinase sequence space to expand the knowledge base on cutinases and their associated PET hydrolase activities. To achieve this, we applied a bioinformatics-driven strategy to select representative sequences of all available members of the bacterial canonical-fold cutinases (PF12695 or IPR029059 family) and several members of the less-studied bacterial minimal-fold cutinases (PF01083 or IPR000675 family). To gain untargeted experimental access to a large number of sequences, we used a gene library, which has been created using Dropsynth technology, combined with a high-throughput screening strategy. For enzyme expression and secretion, we selected *B. subtilis* as host organism, which is highly relevant for industrial enzyme production. This approach enabled the investigation of a library comprising 1,120 DNA sequences encoding 954 unique bacterial cutinases for expression in an industrially relevant bacterial host, and activity against two synthetic substrates, Impranil DLN and semicrystalline PET

## Results and Discussion

### Library showed high coverage and strong bias

Figure 1 shows the distribution of sequences included in the DNA library compared to the Plastics-Active Enzymes database (PAZy) (10) or PETase Activity Natural Dataset (PANDA) database (20). Besides covering a broad sequence space, we also aimed at identifying PET hydrolases within farther phylogenetic clusters, and therefore, we have selected among more distantly related canonical- and minimal-fold cutinases with lower sequence similarity to known enzymes (SI-Figure 1). In total, we have selected 721 canonical-fold cutinases, 221 minimal-fold cutinases, and twelve controls, resulting in 954 unique cutinases in the library. Controls were added as five different DNA sequences using various codons, while canonical- and minimal-fold cutinases were added as one DNA sequence, and one or two DNA sequences, respectively, adding up to 1,120 genes. Pooled sequencing of the raw DNA-library revealed that nearly all sequences were found in the library, but with different distributions. A Gini-coefficient of 0.67 was calculated indicating a strong bias towards certain sequences (SI-Figure 2). Therefore, a Monte-Carlo simulation was performed to estimate the number of unique sequences based on random picking (SI-Figure 3). Based on those results, screening was limited to a colony number expected to maximize sequence diversity while remaining experimentally manageable. We estimated that picking approximately 2,600 random colonies would yield around 600 unique sequences, while increasing the number of colonies would only marginally increase the chances of discovering additional unique sequences.

**Figure 1.**
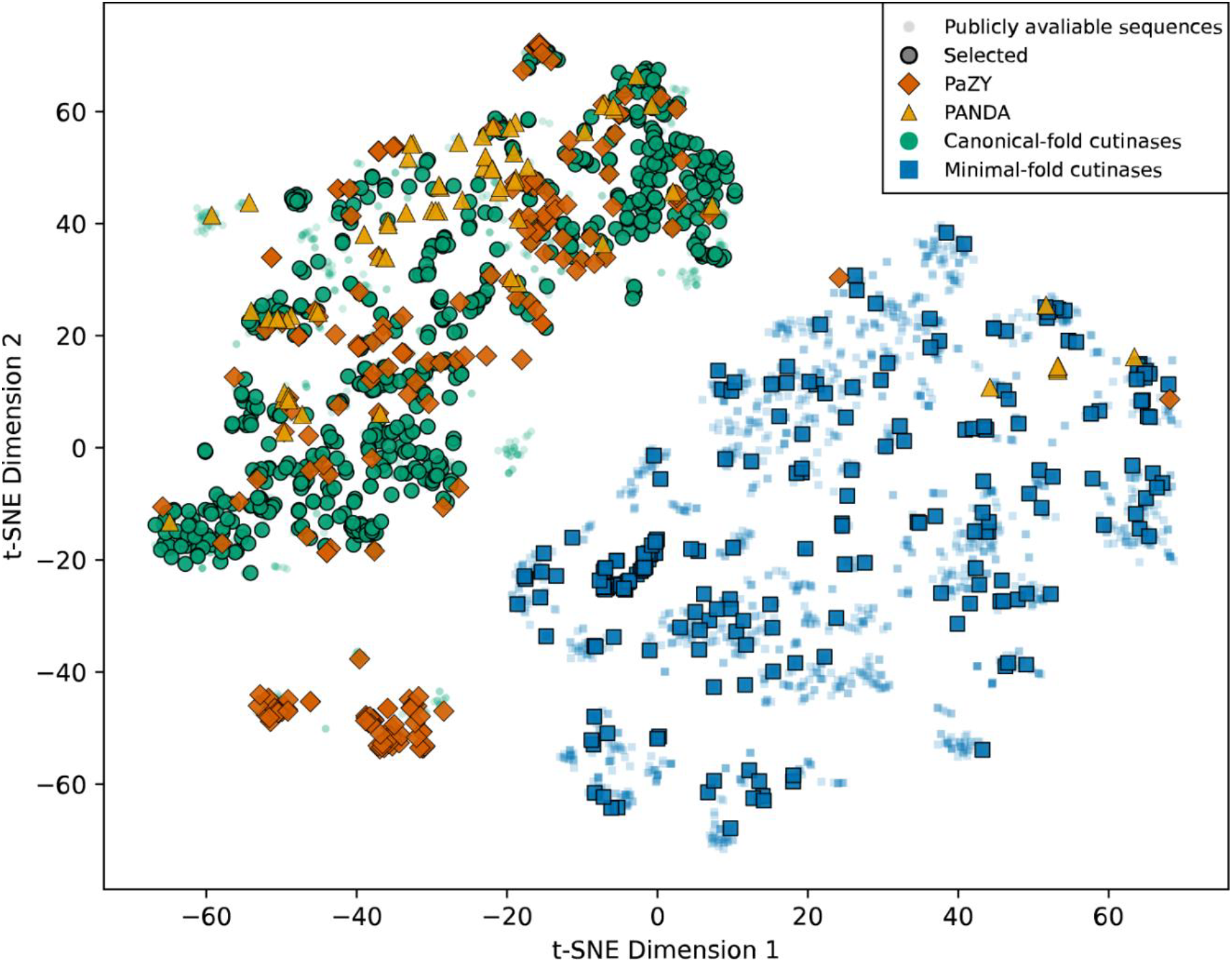
T-distributed stochastic neighbor embedding (t-SNE) visualization of publicly available cutinases at the time of the study. Clustering is based on esm1b-embedding. Sequences selected for cloning and analysis are shown with borders. Canonical-fold cutinases are shown as green dots, minimal-fold cutinases are shown as blue squares, and references with reported activity against PET are shown as dark orange diamonds (PAZy database) or bright orange triangles (PANDA database). Light blue and green dots in the background represent sequences not selected for the library but present in the sequence space.

### Impranil-DLN is a suitable substrate for deselecting inactive cutinases

To establish an efficient screening strategy for excluding inactive or poorly expressed cutinases, we first evaluated a subset of 384 randomly selected clones for esterase and polyester-degrading activity using three substrates: the chromogenic ester substrate p-nitrophenyl butyrate (pNP-butyrate), the aqueous polyester-polyurethane dispersion Impranil DLN, and semicrystalline PET powder. The subset included 32 clone-controls not selected from the DNA library and 352 clones from the DNA Library, which contains replicates as the clones were not sequences before this initial test. Among the 384 clones tested, 28 displayed activity on PET, 279 on Impranil DLN, and 382 on pNP-butyrate. Comparative analysis revealed no significant or very weak correlations (Spearman, SI-Figure 4) between activity on the three substrates, suggesting that ester hydrolysis on simple model substrates does not predict polyester degradation capacity. Nevertheless, all enzymes that were active on PET also exhibited activity on both Impranil DLN and pNP-butyrate. This indicates that these substrates can serve as pre-screening filters for narrowing down candidates for PET hydrolysis. Between the two, Impranil DLN yielded fewer positive hits and thereby reduced the number of false positives carried into subsequent PET assays. In addition, since pNP-butyrate can also easily be hydrolyzed by native *B. subtilis* esterases, it is a less suited screening substrate due to the higher background activity.

### Activity against Impranil-DLN is widely distributed across the sequence space

From the 2,640 colonies screened, we identified 156 unique cutinases with detectable activity against Impranil DLN. Of these, 126 sequences belonged to the canonical-fold cutinases, while 20 sequences belonged to the minimal-fold cutinases (Supplementary Dataset 1) and ten were controls. This distribution closely mirrored the relative representation of the two families in the DNA library (Figure 2, A), suggesting that activity was not biased toward a particular lineage but reflected the overall diversity of the input library.

**Figure 2.**
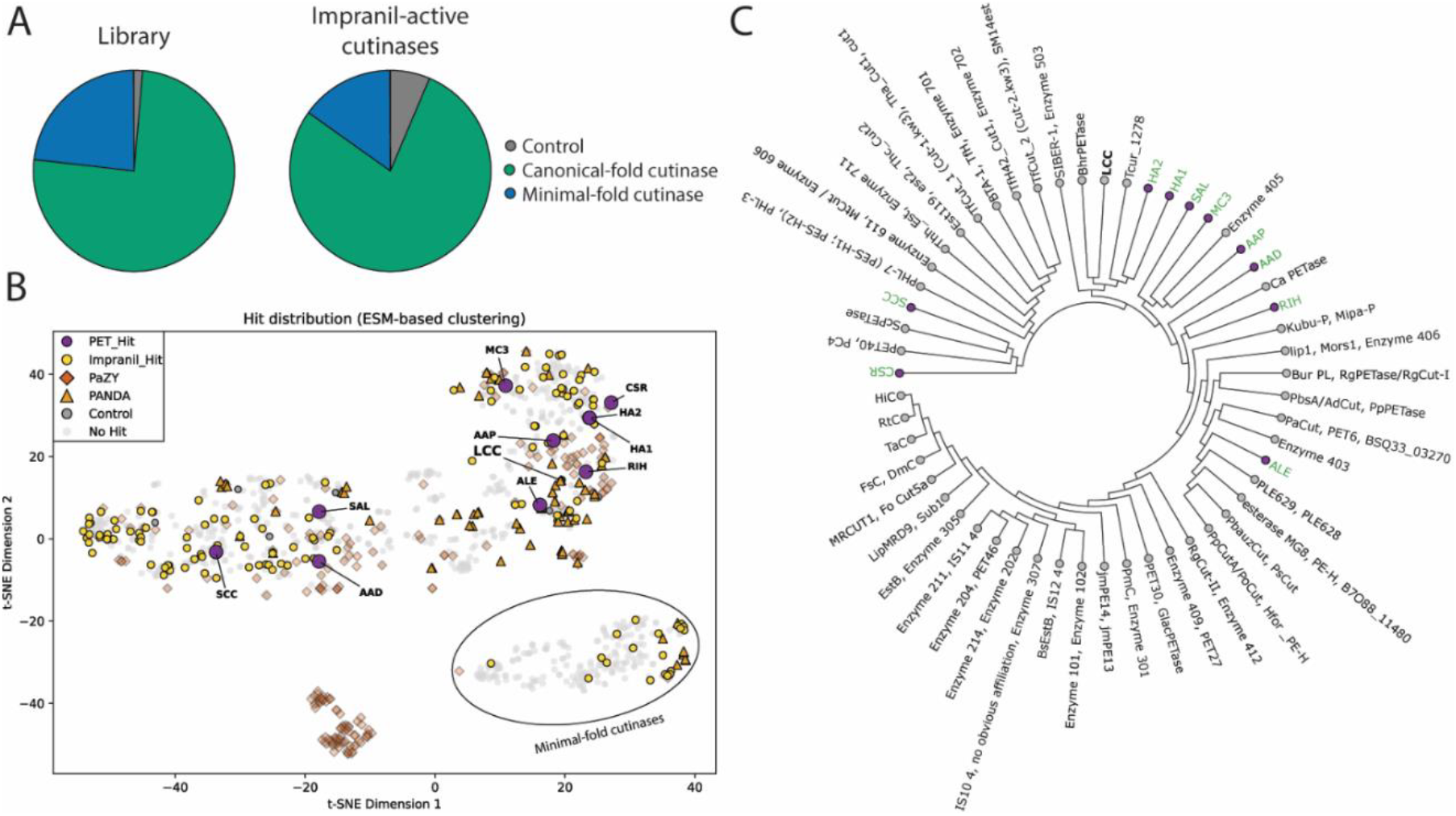
(A) Distribution of cutinases in the library compared to distribution of cutinases with activity against Impranil DLN. (B) t-distributed stochastic neighbor embedding (t-SNE) visualization of esm-1b based clustering of the sequences selected for the library and their respective activities. Cutinases with activity against PET are shown as purple circles with labels. Cutinases identified in this study showing activity against Impranil DLN are shown as yellow circles. Selected cutinases without activity are shown in the background as grey transparent dots. Reference cutinases with reported PET-degrading activity are shown as follows: PAZy database as red diamonds or PANDA database as orange triangles. Control cutinases are shown as grey dots, while LCC is highlighted separately. (C) Phylogenetic relationship between cutinases with activity against PET, and references from the PANDA database. Newly identified cutinases from this work are highlighted in green.

The number of active cutinases was less than expected from the Monte-Carlo simulation, which reflect differences in expression and secretion efficiency across the diverse enzyme set, as all enzymes were screened using a single standardized expression setup without individual optimization.

A notable finding was the successful heterologous expression of 20 bacterial cutinases belonging to the minimal-fold cutinases. Aside from our study, only few other bacterial cutinases from the minimal-fold cutinases have been shown to exhibit activity against unnatural substrates (13, 14). This enzyme class comprises several thousand members, of which fungal homologs have been shown to have activity against PET (21, 22). Therefore, exploring this untapped sequence space holds potential for identifying new enzyme activities. This finding underscores the potential of expanding bioprospecting efforts to enzyme families different from those frequently associated with plastic degradation.

Interestingly, cutinases with activity against ester-based plastics were broadly distributed across sequence space. Using t-distributed stochastic neighbor embedding (t-SNE), we observed that sequences with Impranil DLN activity were not clustered within a narrow clade but rather spread across multiple regions of the diversity landscape (Figure 2,B). Furthermore, it seems that many reference enzymes, listed in the PAZy or PANDA database, form distinct clusters distant from our screening hits, here shown based on structural clustering. This indicates that we identified cutinases outside of the well covered sequence space of PET hydrolases.

### A minor fraction of cutinases is active against PET

Among the identified 146 Impranil DLN-active cutinases (126 canonical-fold, 20 minimal-fold), ten were found to have activity against PET (Table 1). All PET-active enzymes belonged to the canonical-fold cutinases, underscoring the importance of this family in PET hydrolysis. It is worth noting that we applied a quite strict cutoff for designating a screening hit as PET hydrolase compared to other studies. For instance, Norton-Baker et al. used a cutoff of 0.05 increase in absorption of aromatics at 260 nm after 48 h, while we used an increase of 0.2 after 24 h (15). The PET hydrolases identified in this study are closely related to the already reported PET hydrolases, such as LCC or PHL7 (Figure 2, C). This observation again highlights the importance of this sequence space for identifying PET hydrolases.

**Table 1.** Canonical-fold cutinases with activity against PET.

| 3-Letter Code | Organism / Origin | Accession Code |
| --- | --- | --- |
| MC3 | Mock community 3 | WP_289009985 |
| AAD | *Actinomadura atramentaria* DSM 43919 | WP_019633337 |
| CSR | *Cellulomonas* sp. Root485 | TREMBL:A0A0Q7MBH6 |
| HA1 | *Hamadaea* sp. 1 | TREMBL:A0A8J8H438 |
| HA2 | *Hamadaea* sp. 2 | WP_290858125 |
| RIH | *Rhizocola hellebore* | TREMBL:A0A8J3Q2S4 |
| SAL | *Spirillospora albida* | WP_157610457 |
| AAP | *Actinomadura physcomitrii* | TREMBL:A0A6I4MG66 |
| ALE | *Alkalilimnicola ehrlichii* | TREMBL:A0A3E0WVY1 |
| SCC | *Saccharothrix* sp. | TREMBL:A0A1Q4YQI4 |

In this dataset, the comparably low frequency of cutinases with activity against PET is noteworthy. In previous studies, where significant efforts were made to preselect promising cutinase candidates, higher fractions of tested cutinases had activity against PET (15, 16, 23, 24). This implies that the meticulous in-silico preselection distorts the actual representation of activity against PET in this enzyme family. Here we present data suggesting that activity against PET is in fact a rare trait among cutinases. While we find PET hydrolases among canonical-fold cutinases with activity against Impranil DLN with approximately 8% frequency, this frequency appears to be lower in the minimal-fold cutinases, where we could not identify a single PET hydrolase among minimal-fold 20 enzymes active on Impranil DLN.

### PET hydrolases have remarkably similar properties

The ten PET hydrolases identified from the screening were purified and biochemically characterized. The characterization included two known bacterial PET hydrolases, Leaf branch compost cutinase (LCC) and *Thermobifida fusca* cutinase (TfC) for comparison. Overall, at 40°C, the new PET hydrolases showed comparable or higher activity than TfC (Figure 3, A). However, under the current experimental conditions, two of the PET hydrolases, RIH and SCC, even exceeded the activity of LCC, showing 1.8- and 1.6-fold higher activity, respectively.

**Figure 3.**
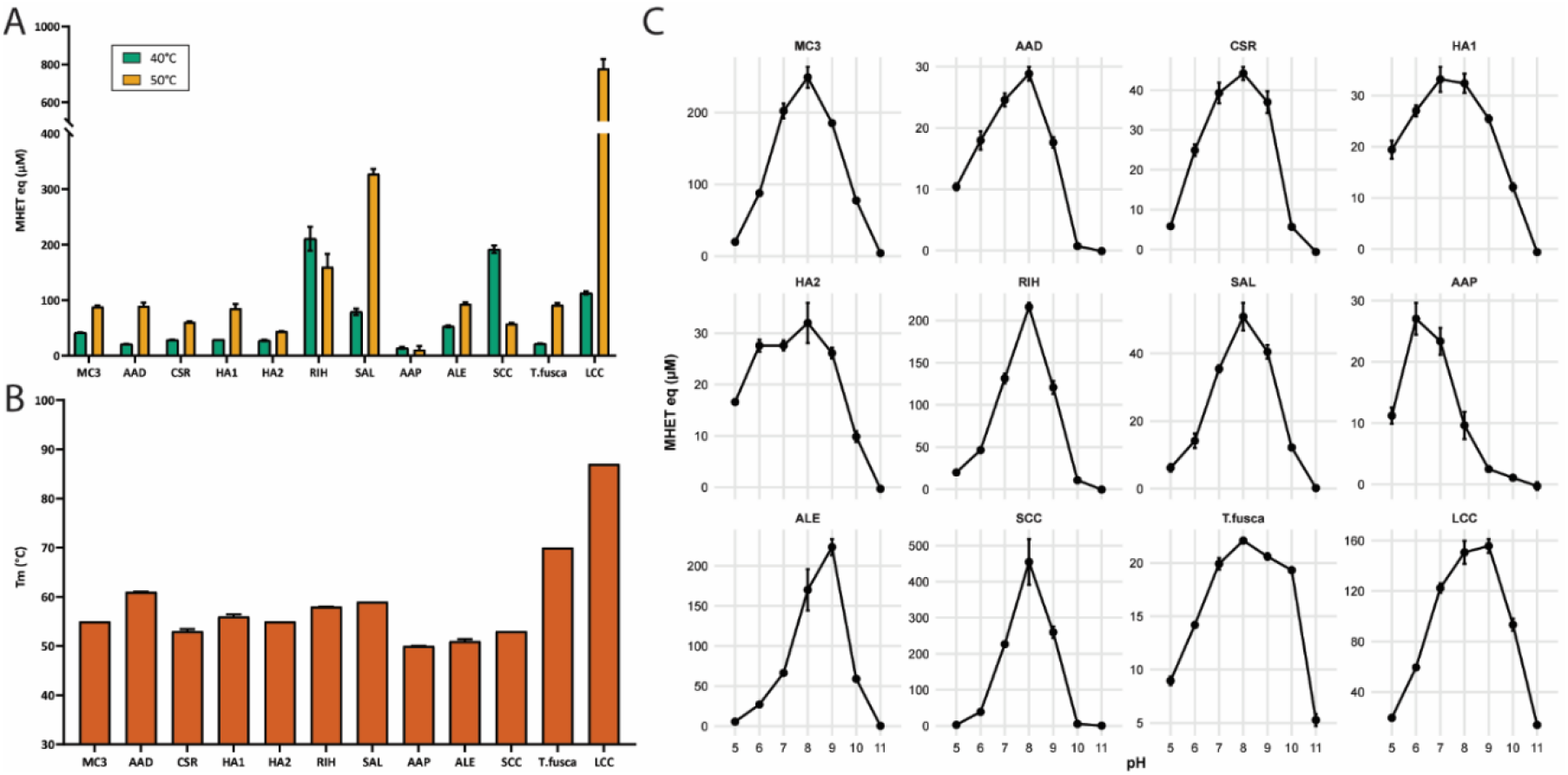
Activity of PET hydrolases in comparison to T. fusca cutinase and LCC. A: MHET equivalents (eq) in µM after incubation for 3 h at 40°C (green) and 50°C (yellow) with enzyme concentrations of 0.1 µM. B: Melting temperature (Tm). C: MHET equivalents (eq) in µM after incubation for 4 h at 40°C with enzyme concentrations of 0.1 µM at pH 5, 6, 7, 8, 9, 10, and 11. Data are mean values ± SD from technical duplicates of the same enzyme preparation.

The activity of the PET hydrolases was also evaluated at 50°C, where all remained active and many gained more activity, with RIH and SCC being the exceptions (Figure 3, A). Compared to TfC, the PET hydrolases SAL and RIH showed 3.5- and 1.7-fold higher activity, respectively, while the remaining enzymes had comparable or lower activity. At 50°C, none of the new PET hydrolases exceeded the activity of LCC. This finding is consistent with the measured denaturation temperatures of the PET hydrolases, which range from 50 °C to 60 °C as determined by nanoDSF, explaining their reduced stability and activity at higher reaction temperatures. Even though RIH and SAL show similar denaturation temperatures, the activity for RIH drops with increasing temperature, while the activity of SAL increases. A similar effect can be observed for other enzymes such as ALE and SCC as well, indicating that other factors than the melting temperature play a critical role in influencing the behavior of PET hydrolases at different temperatures.

Furthermore, the pH dependence of the new PET hydrolases was examined at 40°C given the industrial relevance of acid tolerant PET hydrolases, as the release of terephthalic acid during PET hydrolysis leads to acidification, necessitating base addition and thereby increasing process costs (7). The PET hydrolases in general exhibited optimal activity near pH 8, which is typical of reported PET hydrolases. Our screening was conducted at pH 8.0, which favored the identification of cutinases with pH optima in the neutral to mild alkaline range. However, one enzyme, AAP, showed maximal activity at pH 6. Interestingly, enzymes MC3, RIH, and SCC displayed higher activity than LCC below pH 8. HA1 and HA2 also showed comparably high activity across a broad range of pH (pH 5-10), comparable to TfC, while most of the other enzymes showed activity at pH between 6-9. The slight discrepancies between activities could potentially be explained by the different reaction conditions between the experiments.

### Conclusions

We presented a systematic and unbiased approach towards exploring the bacterial cutinase sequence space with respect to activity against ester-based plastics. The combination of computational discovery for candidate selection, multiplexed gene synthesis, and high-throughput screening offers a scalable strategy for expanding the diversity of PET hydrolases. Here, we screened a library of cutinases, representing both the canonical-fold cutinases and the minimal-fold cutinases. From 2,640 clones, we identified 156 unique enzymes including ten controls, expressed and secreted in *B. subtilis* with activity against a synthetic polyester. Those enzymes were broadly distributed across the cutinase sequence space. Among these, we discovered ten PET-active cutinases, all belonging to the canonical-fold cutinases. These findings extend the diversity of known PET hydrolases into phylogenetically distant clusters so far underrepresented in the PAZy or PANDA databases. While 20 minimal-fold cutinases displayed activity on the polyester substrate Impranil DLN, none showed detectable activity against PET. Up to now, only two bacterial enzymes of this family have been studied for their ability to degrade synthetic polymers (13, 14). Therefore, our findings expand the functional diversity of this enzyme class towards plastics.

The biochemical characterization revealed that the ten PET-active cutinases shared the alkaline activity optima typical of known PET hydrolases, but several enzymes retained activity at slightly lower pH, which is an advantageous trait under industrially relevant conditions where terephthalic acid release causes acidification. Several of the newly discovered enzymes showed compatible or higher activity than LCC at lower temperatures. However, due to their modest thermostability (Tm 50-60°C) and thus limited performance at elevated temperatures, none surpassed the benchmark enzyme LCC in high-temperature PET depolymerization.

Together, these results demonstrate the feasibility and power of systematic, large-scale exploration of enzyme sequence space for discovering novel polyester hydrolases. Our findings complement other recent studies, which aimed at experimentally exploring unknown sequences, where most enzymes showed low to moderate activities (15, 16, 20, 23, 24). Considering that we have identified 146 cutinases with activity against Impranil DLN, and only ten of those showed activity against PET, none with exceptional stability or activity, we conclude that high PET activity among cutinases is a rare trait among canonical-fold cutinases. Furthermore, the activity levels detected of the cutinases remained at remarkably similar levels. Our study also highlights that the ability to hydrolyze synthetic polyesters is not restricted to a small number of enzymes with special features, but rather a property broadly distributed across cutinase families and across structure and sequence space.

Our findings raise the question about the origin of the highly active PET hydrolases such as LCC, BhrPETase, or PHL7, or more recently Mipa-P or Kubu-P, which so far remained hidden gems within the sequence space (9, 16, 25, 26). So far, no clear explanation of their high activities and biological relevance was provided, and only targeted enzyme mining has resulted in identification of remarkably similar enzymes. For future studies, we suggest broadening the scope of enzyme discovery towards exploring more distant phylogenetic clusters of thermophilic lineages in an unbiased manner to better understand the nature and origin of those enzymes.

## Materials and Methods

### Gene selection and library design

All available bacterial sequences from the two enzyme families, canonical-fold cutinases (PF12695 or IPR029059 family) and minimal-fold cutinases (PF01083 or IPR000675 family), were clustered and analyzed based on mature protein similarity to ensure coverage across both well-characterized PET-active clades and more divergent branches. We excluded sequences originating from potential pathogenic organisms (Risk Group 2) and applied a length restriction of minimum 475 bp and maximum 1000 bp to stay within the requirements for Dropsynth technology. The sequences were embedded using the esm1b protein language model (esm1b*t33*650M_UR50S, available at the time of the study). In parallel, a sequence-based representation was derived using a k-mer (3-mer) approach, followed by L2-normalization and dimensionality reduction using TruncatedSVD. Clustering in both embedding spaces was performed using DBSCAN with a cosine distance metric. For visualization, dimensionality reduction was performed using t-distributed stochastic neighbor embedding (t-SNE). Based on this embedding, representative genes were selected. We included a selection of relevant known PET hydrolases as controls (SI-Table 1). No further enrichment for thermostability or other properties was done. To facilitate cloning, each sequence was designed to contain restriction enzyme recognition sites. To facilitate the DNA assembly by Dropsynth technology, random sequences were added to the genes, so that all have the same length.

### Transformation, colony picking, and expression

Based on a service agreement between Novonesis and Calin Plesa, the DNA library was synthesized as previously described using Dropsynth technology (17). After delivery, the library was cloned into an *E. coli* vector using restriction enzyme-based cloning and transformed into *E. coli* Dh5α cells for long term preservation. At this stage, the pooled library was sequenced for quality control and bias determination. All cloning procedures were done using standard molecular biology techniques as described previously (27, 28). Transformed *B. subtilis* was plated on LB-agar containing 6 µg/ml chloramphenicol and a total of 2,640 transformed colonies were picked randomly into 96-well microtiter plates containing 200 µl LB-medium with 6 µg/ml chloramphenicol. The plates were incubated at 37°C overnight and supplemented with glycerol to a final concentration of 15% and stored at -80°C.

For screening of clones, precultures were prepared by inoculating 150 µL TB-glycerol medium containing 6 µg/ml chloramphenicol with 5 µL of colony culture, followed by incubation at 37°C and 700 rpm overnight. Expression was performed in 96-deep-well plates (Nunc Polypropylene DeepWell™, Thermo Fisher Scientific), by inoculating 600 µL expression medium with 8 µL preculture. Plates were incubated at 30°C, 700 rpm for 3 days, followed by centrifugation at 3000 x g for 20 minutes to harvest expression supernatant for screening.

Shake flask expression was performed for purification of the PET active cutinases. Precultures were prepared by inoculating 6 mL TB-glycerol medium containing 6 µg/ml chloramphenicol with transformants cloned with PET active cutinases in 50 mL tubes (Nunc, Thermo Scientific), and incubated at 37°C, 700 rpm overnight. Protein expression was carried out in 500 mL baffled shake flasks containing 150 mL expression medium inoculated with 2 mL preculture. Flasks were incubated at 30°C, 270 rpm for 3 days. After incubation, 45 mL of the expression broth was transferred to 50 mL centrifugation tubes centrifuged at 14000 x g, 10°C for 30 min. The resulting supernatant was filtered (Whatman, 0.45 μm) and collected for purification. Prior to purification, the pH of 35 mL filtered supernatant was adjusted to pH 7.4, and imidazole was added to a final concentration of 20 mM.

### Activity Assays for Supernatant Screening

Activity on pNP-butyrate was evaluated in 100 µL 250 mM sodium-phosphate buffer, pH 8, by adding 1% (v/v) expression supernatant to a 0.1% (v/v) pNP-butyrate solution in microtiter plates. Plates were incubated at room temperature for 3 minutes and the increase in absorbance was measured at 405nm. Clones that resulted in an Abs405 increase of 0.1 or more were considered active on pNP-butyrate.

For detecting activity against Impranil DLN (Covestro), a 1% suspension of Impranil DLN was prepared in 100 mM Universal Buffer 2 (UB2) (200 mM Tris-HCl, 200 mM Bis-Tris, and 200 mM sodium acetate) (29). Expression supernatant was added to a final volume fraction of 5% in microtiter plates containing the Impranil DLN suspension. Plates were incubated at 30°C for 2 h, and the decrease in turbidity was quantified spectrophotometrically. Clones that resulted in a decrease in Abs600 of ≥ 4% were considered active against Impranil DLN.

For experiments with PET, a semi crystalline PET powder purchased from Goodfellow Co. UK with crystallinity > 50% and typical particle size of around 300 µm was used. Screening on PET was performed in microtiter plates using a PET load of 10 g/L suspended in 250 mM sodium phosphate buffer, pH 8, and expression supernatant volume fractions of 2.5% and 5%. Reactions on PET were performed at 40°C for 24 h, followed by centrifugation at 3000 x g. Subsequently, 100 μL of supernatant was analyzed at 240 nm to quantify the absorbance of released soluble aromatic products as a measure of activity (30). Clones showing an increase in absorbance at 240 nm of ≥ 0.2 were considered active on PET.

### Purification

PET active cutinases were purified using the His GraviTrap^TM^(Cytiva 11-0033-99) system according to the manufacturer’s instructions. All fractions were collected and analyzed using SDS-PAGE.

The buffer of the eluate was exchanged to a 100 mM HEPES pH 7.65 buffer using a PD10 desalting column, Sephadex G-25 M (Cytiva 17-0851-01). The desalting column was equilibrated by applying 30 mL 100 mM HEPES buffer, pH 7.65, before adding 2.5 mL of the eluate to the column. To release the protein, 3 mL 100 mM HEPES buffer, pH 7.65, was added to the column and the purified protein was collected.

*Thermobifida fusca* cutinase (TfC) and leaf-branch compost cutinase (LCC) were produced according to published protocols (28, 31, 32).

### Biochemical characterization of PET active enzymes

The thermal transition midpoint (T_m_) of the PET active enzymes was determined by nano differential scanning fluorimetry (nanoDSF). Enzyme concentration was adjusted to 4 µM in 100 mM sodium phosphate buffer (pH 8) and analyzed in duplicates using standard Prometheus capillaries on the Prometheus Panta (NanoTemper) instrument at 100% excitation power using a temperature ramp of 2°C/min from 25°C-95°C.

PET degrading activity was assessed in duplicate in non-binding 96-well microtiter plates (Greiner Bio-One) using a PET load of 10 g/L suspended in buffer, an enzyme concentration of 0.1 µM, and a final reaction volume of 200 µL. Plates were incubated in a thermomixer (Eppendorf) at 1100 rpm. Reactions were centrifuged at 3000 × g for 3 min, and 100 µL of supernatant was analyzed spectrophotometrically at 240 nm. Activity was reported as MHET equivalents as described elsewhere (30). For temperature dependency assays, reactions were performed in 50 mM sodium phosphate buffer (pH 8) and incubated at 40 °C or 50 °C for 3 h. For pH dependency assays, reactions were performed in 50 mM Britton–Robinson buffer (33) with fixed ionic strength of 0.2 mM at pH 5–11 and incubated at 40 °C for 4 h.

### Statistics and Reproducibility

The number of colonies screened was selected based on a Monte Carlo simulation estimating the number of unique sequences recovered by random colony picking. Screening data from supernatants was obtained from randomly picked colonies without deliberate replication. Biochemical characterization of purified enzymes was performed in duplicates and data is presented as mean ± standard deviation (SD). Pearson and Spearman correlation analyses were used to assess associations between activities measured on PET, Impranil DLN, and pNP-butyrate for 384 clones. Technical replicates refer to repeated measurements of the same enzyme preparation.

If your research involved human or animal participants, please identify the institutional review board and/or licensing committee that approved the experiments. Please also include a brief description of your informed consent procure if your experiments involved human participants.

## Acknowledgments

This work was supported by Innovation Fund Denmark (grant no. 0224-00033B) and Novonesis A/S.

