## Supporting Information for "Mapping bacterial cutinase sequence space by high-throughput screening reveals that PET hydrolysis is a rare property"

### Supplementary Figures

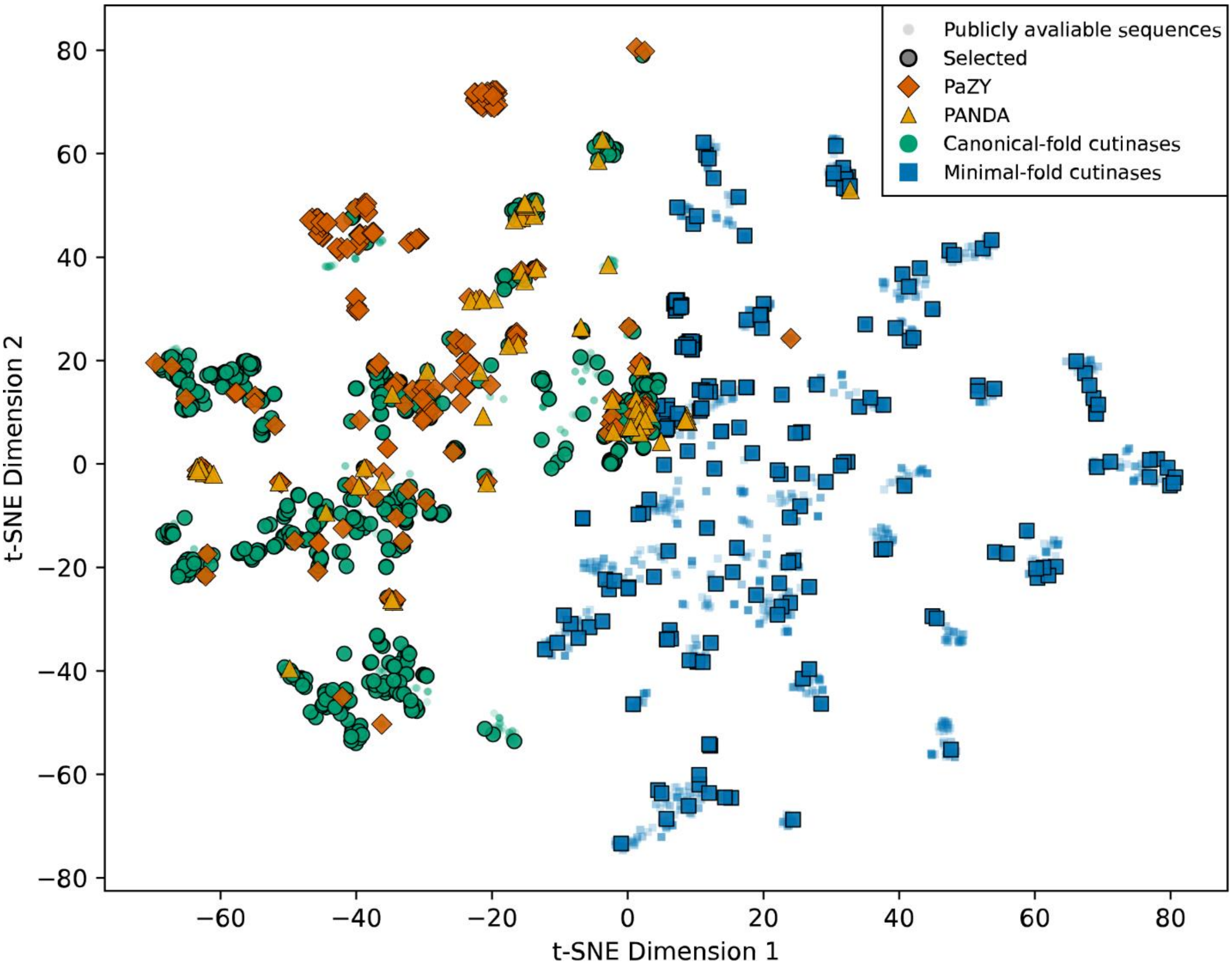


**Figure S1**. T-distributed stochastic neighbor embedding (t-SNE) visualization of publicly available cutinases at the time of the study. Clustering is based on sequence similarity. Sequences selected for cloning and analysis are shown with borders. Canonical-fold cutinases are shown as green dots, minimal-fold cutinases are shown as blue squares, and references with reported activity against PET are shown as dark orange diamonds (PAZy database) or bright orange triangles (PANDA database). Dots in the background represent sequences not selected for the library.

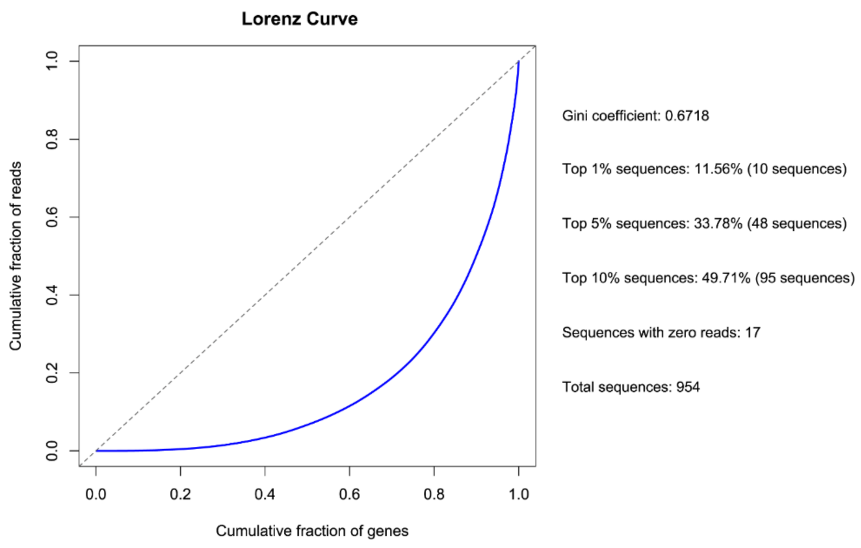


**Figure S2**. Cumulative fraction of sequencing reads against cumulative fraction of protein sequences, showing that a small number of sequences have high abundance, while others appear only in very few copies.

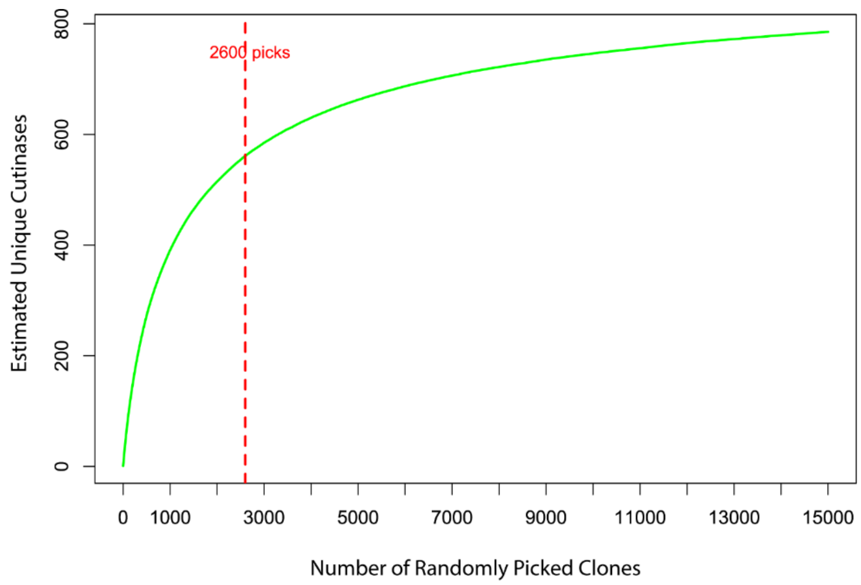


**Figure S3**. Simulated random picking from the cutinase library using a Monte-Carlo simulation based on sequencing reads. Number of randomly picked clones is plotted against the number of estimated unique cutinases. We have picked approximately 2,600 clones, which is indicated on the graph as red-dotted line.

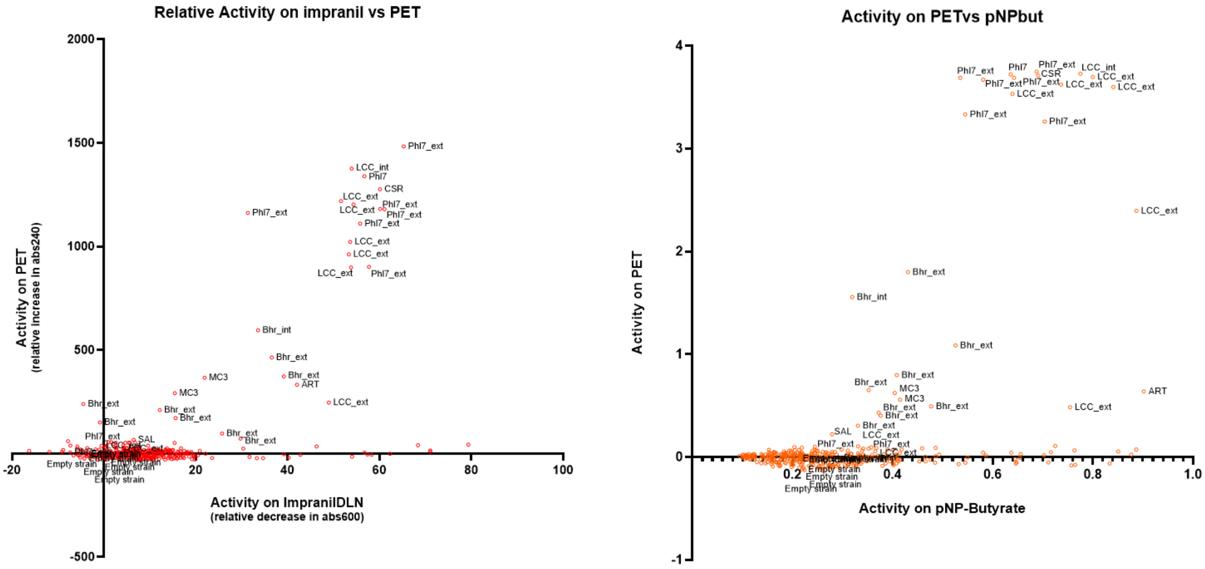


**Figure S4**. Correlation between activity against PET and activity against Impranil DLN (left panel) and pNP-butyrate (pNPB, right panel). The following correlations were calculated (n = 384): Pearson: PET vs pNPB: r = 0.49, p < 0.0001, PET vs Impranil: r = 0.58, p < 0.0001, pNPB vs Impranil: r = 0.41, p < 0.0001. Spearman: PET vs pNPB: r = 0.12, p = 0.0214, PET vs Impranil: r = 0.10, p = 0.0621, pNPB vs Impranil: r = 0.04, p = 0.3940.

**Supplementary Tables**

**Table S1**. Controls used in the DNA library. Cutinases with reported activity against PET.

| Accession Code | Description | Abbreviation | Reference |
|---|---|---|---|
| G9BY57 | Leaf-compost cutinase | LCC | [1] |
| PDB:7NEI | Polyester Hydrolase Leipzig 7 | PHL7 | [2] |
| A0A0K8P6T7 | *Ideonella sakaiensis* | IsPETase | [3] |
| A0A0G3BI90 | *Polyangium brachysporum* | PET12 | [4] |
| A0A1M5F0K3 | *Vibrio gazogenes* DSM 21264 = NBRC 103151 | PET6 | [4,5] |
| A0A2H5Z9R5 | Bacterium HR29 | BhrPETase | [6] |
| C3RYL0 | Uncultured bacterium | PET2 | [4] |
| D1A2H1 | *Thermomonospora curvata* DSM 43183 | Tcur0390 | [7] |
| D1A9G5 | *Thermomonospora curvata* DSM 43183 | Tcur1278 | [7] |
| Q47RJ6 | *Thermobifida fusca* YX | Tfu_0882 | [8] |
| R4YKL9 | *Oleispira antarctica* RB-8 | PET5 | [7] |
| W0TJ64 | *Saccharomonospora viridis* | Cut190 | [9] |
